# The cognitive mechanisms of state transformation during the multi-state maintenance of visual working memory

**DOI:** 10.1101/2022.04.29.490000

**Authors:** Ziyuan Li, Qiang Liu

## Abstract

Visual working memory (VWM) system serves as a cornerstone for high-level cognition. The state-based models of working memory delineated a hierarchy of functional states. Memory representations in the passive state are robustly maintained while the active representations were effectively processed at the same time. The memory representations are dynamically transferred between the two states according to task demands; however, it was still unclear how the state transformation process implemented to achieve a perfect storage-dissociation. To explore the property of transformation process, we adopted a sequential presentation paradigm where two memory arrays were presented sequentially; that effectively directs memory items for retention in the two distinct states. We modulated the temporal context concerned with the state transformation process, presentation time of the second array and retention interval between memory arrays. These results indicated that state transformation reflected a consolidation process of memory representations from the active into the passive state; this transformation process is subject to cognitive control. Moreover, we found that sufficient temporal context facilitated a smooth state transformation, thus minimizing the memory loss. These findings lead to a further understanding for the storage mechanism of working memory representations during the dynamic processing.

## Introduction

Visual working memory (VWM) plays a foundational role in advanced cognitive functions. It serves to retain and manipulate the information in mind (Aben et al., 2012; Baddeley, 1992; Eriksson et al., 2015; Miller et al., 2018). The relevance of representations to the current task and the activation level of memory representations have been described by the state-based theory (de Vries et al., 2020; Nee & Jonides, 2013; Oberauer, 2002; LaRocque et al., 2014). Memory representations with great attentional priority are presumed to be retained in an active state, dependent on recurrent neural activity (Lewis-Peacock et al., 2012; Manohar et al., 2019; Stokes, 2015). Whereas those receiving less priority are relegated to a passive state (i.e., latent state or activity-silent state) for later use; these passive representations received robust maintenance in a silent manner which was underpinned by the changes of short-term synaptic plasticity (Mongillo et al., 2008; Kamiński & Rutishauser, 2020; Muhle-Karbe et al., 2021). During the maintenance of working memory representations, the passive representations are mutually dissociated from the online processing of active representations (Li et al., 2021; Oberauer, 2002).

The view that memory representations are dynamic and modifiable is supported by the observation that passive representations can be reactivated by application of transcranial magnetic stimulation (TMS) or a visual impulse stimulus (Rose et al., 2016; Wolff et al., 2015, 2017). Accumulating evidence has since confirmed that the passive state can provide genuine maintenance for memory representations, while also protecting them from interference and time decay (Johanna Kreither, 2021; Stokes et al., 2020; Zhang et al., 2022). Thus, transferring the prospectively relevant items into the passive state for temporary maintenance seems to be an ecologically optimal way to satisfy the cognitive requirement of processing dynamically presented visual information (Rose, 2020; Stokes, 2015).

The state transformation contributes to the functionally different roles of the memory representations in the active and passive states (LaRocque et al., 2014; Masse et al., 2020; Olivers et al., 2011; Stokes et al., 2020). However, there was rare work to explore how the state transformation of memory representations is implemented during the memory maintenance. The investigation on the mechanisms of state transformation promoted us to understand the dynamic coding framework. According to the proposal that the active and passive states were underlain by different neural modes, it could be presumed that the state transformation possibly signified the formation of synaptic plasticity after the abolishment of persistent neural activity. This implied that the to-be-transferred representations would undergo a consolidation process into the passive state via state transformation, which might be cognitively resource-consuming. An alternative presumption was that the state transformation into the passive state resulted from the attenuation of persistent neural activity, due to the memory representations were proposed to be maintained in the combination of persistent activity and short-term synaptic plasticity (Masse et al., 2020; Mongillo et al., 2008). Such that the state transformation did not involve cognitive consolidation. These presumptions inspired us to explore whether the state transformation mirrored a consolidation process of memory representations into the passive state.

Prior research provided us an effective experimental paradigm to unveil the question. In Experiment 3 of Zhang et al.’s research, the EEG results revealed that the first presented array was successfully transferred to the passive state during the active processing of the second array, which was manifested by the dropout of CDA component from the first to the second delay (Zhang et al., 2022). Their behavioral results pattern showed an active load effect on the passive representations, which, however, was quite incompatible with the resource-dissociation account, in that this account posited a clear dissociation between the active memory processing and passive memory maintenance (Li et al., 2021). The incompatibility could be interpreted by an overlap between the preceding state transformation and subsequent active processing due to a tight temporal context, if the state transformation was equivalent to a consolidation process of initially presented items into the passive state. In such case, the active load effect could be avoided by extending the presentation of the second array, because the state transformation could consolidate the to-be-transferred representations into the passive state prior to the subsequent active processing. In contrast, if the state transformation manifested the spontaneous attenuation of persistent activity, the incompatibility with resource-dissociation account could be accounted for by the explanation that the activity pattern of the first presented array encountered the subsequent sensory processing. Specifically, the neural activity of initially presented items did not return to the baseline level when new stimuli triggered activity of sensory processing, thus inducing the active load effect on the to-be-transferred representations. This effect was independent of the temporal length of subsequent memory event.

In this study, we attempted to explore the cognitive mechanisms of state transformation by adopting sequential presentation tasks. Basing on the experimental paradigm and parameters of Zhang et al.’s experiments, we first modulated the presentation time of the second array in Experiments 1 and 2. If the state transformation reflected a consolidation process of memory representations into the passive state, we predicted that extending the presentation time of the subsequent array could eliminate the overlap with the subsequent active processing, thus no active load effect on the passive representations. If it was the case, the transformation process would always overlap with subsequent active processing when the temporal context was beyond participants’ anticipation, because the uncertainty of temporal context placed urgent demands of consolidating sensory stimuli into the active state. This question was explored in Experiment 2. To further explore the property of state transformation, Experiment 3 modulated the interval retention between two memory arrays, and predicted that the state transformation could smoothly complete before the presentation of sensory stimuli if extending the delay retention sufficiently; this would additionally evidence that the occurrence of state transformation was not necessarily triggered by subsequent stimuli. Overall, these findings demonstrated that the state transformation intrinsically mirrors the consolidation process into the passive state; this process unfolds over time and is cognitively controlled.

## Experiment 1

This experiment sought to explore whether the state transformation reflected the consolidation process of memory representations into the passive state. A sequential change detection task was adopted, where 2 items in memory array 1 were potentially retained in the passive state as they were probed at the end of a trial, while 2 or 4 items in memory array 2 were retained in the active state in that they were immediately probed after a short delay. This allowed us to explore whether active load variation had effect on the passive representations. Ye et al. (2017) proposed that the resources allocation was subject to the exposure duration of stimuli. Their results indicated that when presented with four items, an exposure duration of 200 ms limited the processing to the involuntary phase, whereas an 500 ms-exposure allowed for the entry into the late, voluntary phase, where cognitive processes were controlled (Ye et al., 2017). Likewise, we chose 200 ms vs. 500 ms as presentation time of subsequent stimuli; such that memory items could gain access to processing at 200 ms presentation condition, but subject to cognitively different one at 500 ms presentation condition. We predicted that, if the state transformation mirrored the consolidation process, the two processes of state transformation and subsequent processing would be compressed into a tight space of time, resulting in an overlap of them only in 200 ms presentation condition; whereas no overlap in 500 ms presentation condition. The overlap was manifested by the active load effect on the passive representations.

Whereas if the state transformation reflected a process of activity pattern attenuating naturally, that active load effect on the passive would be consistently observed regardless of the presentation time of subsequent stimuli.

## Method

### Participants

Sample-size determination was based on previous study that indicate the magnitude of the effect could be expected (Li et al., 2021). We used a similar sample size (30 in each experiment) to ensure the findings were reliable. None of the participants overlapped across experiments. Each participant has signed informed consent forms prior to participation, would receive 30 CNY after the completion.

Their vision and color vision were normal or corrected-to-normal. In Experiment 1, 30 normal Chinese college students (six males; mean age: 22.07 ± 2.22 years) were recruited. All experiments complied with the Declaration of Helsinki. The present research has gained permission from the Human Research Institutional Review Board in Liaoning Normal University (number.: 20211010).

### Apparatus and stimuli

We employed E-Prime 2.0 to run the experiment procedures in a 19-inch LCD monitor with refresh rate of 60-Hz (1920 × 1080 pixels). A fixation cross (0.23°) was shown centrally in the screen, never disappeared through the whole trials. Colored squares were selected as memory stimuli (0.49° × 0.49°). The color of memory squares was selected from a color pool (blue, black, red, magenta, lime, purple, yellow, and cyan) without repetition within a trial. The squares in memory array 1 were located horizontally at the level of fixation cross in the left and right sides. In memory array 2, the squares were located along the midline when two items were presented, while they were located in the left and right sides separately with two in each side when remembering four items. The locations of all memory items were consistently on a invisible square (1.80° × 1.80°) around the fixation. In the two probe arrays, the locations previously presenting memory items were occupied by square outlines, one of which was colored. The locations of items in probe 1 corresponded with the memory array 2, and probe 2 corresponded with memory array 1. The distance between participants and screen fixed at 70 cm. The height of fixation cross leveled with participants’ eyes.

### Procedure

As depicted in Figure 1, the first array was presented for 0.2 s for encoding two items, which was followed by a delay interval of 0.8 s. Then the second array appeared for either 0.2 s or 0.5 s in a blocked design; meanwhile, two or four items were randomly presented in each trial with an equal probability. After the second delay, two probe arrays sequentially appeared to detect the memory of memory array 1 and then memory array 2 in a reverse order manner. The two probe arrays were separated by an interval of 1 s. The accuracy was stressed over speed. The presentation of probe arrays thus had no time limit, which disappeared once making response. If the colored items in the probe arrays were the same as the memory items in the corresponding locations, participants made “same” response by pressing “Z” in the keyboard, or making “different” response by pressing “M”. The same and different responses for each probe were fully random with 50% probability in each block. In the different trials, the color of probe items was selected from eight possible colors mentioned above, but never used by previous memory items in one trial.

**Figure 1:**
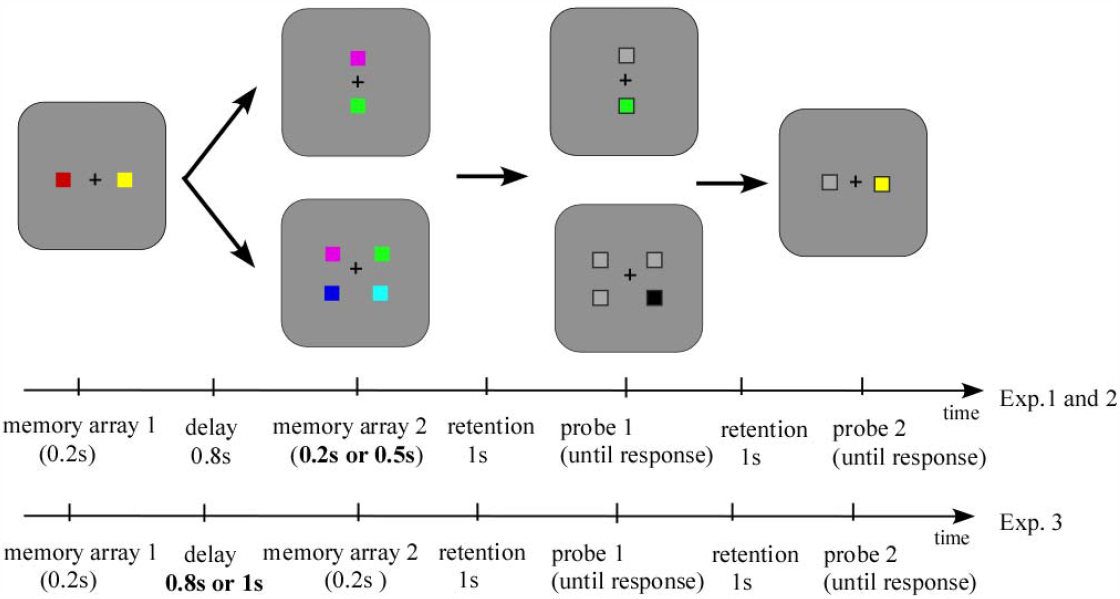
The schematic diagram of the experimental events. The colored squares are designed as memory items. Memory array 1 always presented two items; memory array 2 presented two or four items randomly within a trial. Memory array 1 and memory array 2 were detected by probe 2 and probe 1, respectively. The time frame of memory events is indicated by the numbers below.

Participants need to perform the short and long presentation conditions with five blocks each, creating a total of 320 trials. A half of participants performed the short presentation blocks first and the long presentation blocks later, while the other half performed the long presentation blocks first and the short presentation blocks later. Before the formal trials, we provided an instruction and a practice session of 8 trials to ensure that each participant was familiar with the procedure. To prevent the contamination of verbal encoding, participants were asked to perform the subvocal rehearsal of digits when performing the memory tasks (Shaffer & Shiffrin, 1972).

### Data analysis

Data were analyzed by JASP software across the three experiments (Love et al., 2019). We reported *η*^*2*^*p* to index the effect size. For t-test concerning the load effect on active and passive memories, Cohen’s d was calculated to show effect size. Bayes factors were included to quantify the evidence for the alternative over the null hypothesis (Schmalz et al., 2021).

## Results

Probes 1 and 2 indicated the accuracy of active representations and passive representations, respectively. This experiment was designed in a within-subject manner, including three variables, active load, representational state and presentation time. A 2 (active set size 2 vs. active set size 4) × 2 (active state vs. passive state) × 2 (0.2 s vs. 0.5 s) ANOVA was conducted, and the results showed a significant interaction of presentation time and load, *F* (1, 29) =8.229, *p* = 0.008, *η*^*2*^*p* = 0.221, and the significant interaction between representational state and load, *F* (1, 29) =81.676, *p* < 0.001, *η*^*2*^*p* = 0.738. The participants’ accuracy depending on the short and long presentation conditions is then analyzed by a 2 (active set size 2 vs. active set size 4) × 2 (active state vs. passive state) ANOVA, and depicted in Figure 2.

**Figure 2:**
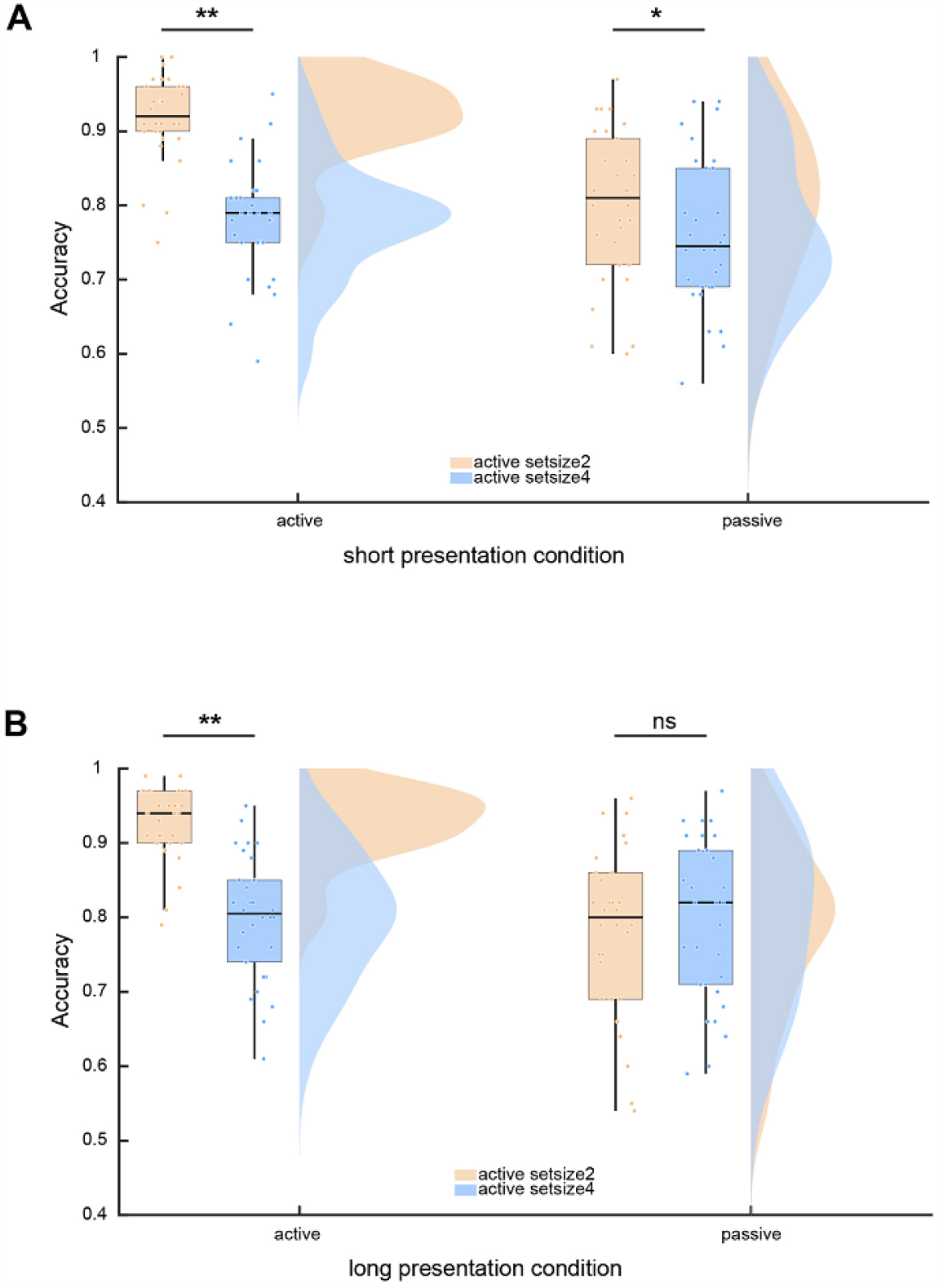
Memory accuracy of active and passive representations in the short (A) and long (B) presentation conditions in Experiment 1. Boxplot, the middle lines denote the median; the box outlines represent the 1^st^ and 3^rd^ quartiles. Scatters indicate data distribution for each condition.

For the short presentation condition (Figure 2A), the 2 × 2 ANOVA results showed a significant interaction, *F* (1, 29) = 35.603, *p* < 0.001, *η*^*2*^*p* = 0.551. That was caused by a larger accuracy decrease for the active memory relative to the passive memory. Subsequent simple effect analysis showed that the active memory accuracy had a significant decrease when the load increased from two to four, *t*(29) = 14.699, *p* < 0.001, Cohen’s *d* = 2.684, *BF10* = 1.115e+12. Also, the passive memory accuracy decreased when increasing the active set size, *t*(29) = 3.658, *p* = 0.001, Cohen’s *d* = 0.668, *BF10* = 32.991. The results pattern showed an active load effect on the passive state in the short presentation condition. While for the long presentation condition (Figure 2B), there was a significant interaction, *F* (1, 29) = 62.279, *p* < 0.001, *η*^*2*^*p* = 0.682. This was driven by a lack of load effect on the passive memories. A subsequent t-test on these data suggested that the active accuracy consistently impaired due to the larger set size, *t*(29) = 8.846, *p* < 0.001, Cohen’s *d* = 1.615, *BF10* = 1.148e+7. However, the passive memory accuracy of was comparable in the conditions of active set size 2 and 4, *t*(29) = 1.121, *p* = 0.272, Cohen’s *d* = 0.205, *BF10* = 0.344. These results suggested that, unlike the short presentation condition, there was no active load effect on the passive representations in the long presentation condition.

## Discussion

These results presented suggested that extending the presentation time of subsequent array effectively eliminated the active load effect on the passive representations. That allowed us to evidence that the state transformation acted as a consolidation process that consolidated the memory representations to the passive state. In such case, sufficient presentation time permitted a smooth state transformation prior to the subsequent sensory processing, thus avoiding an overlap and resource competition. While a tight temporal context drove participants to initiate the subsequent processing with the incompletion of state transformation, thereby causing an overlap. These results pattern denied the hypothesis that the state transformation of memory representations signified just the natural attenuation of persistent activity pattern of memory representations.

Based on these findings, we found that participants’ awareness of temporal context guided them to adopt different strategies corresponding to the short and long temporal contexts. Nevertheless, it should be expected that a conservative strategy was adopted to both the tight and sufficient contexts when the temporal context was unpredictable. That is, the two processes of state transformation and subsequent sensory processing would always be compressed into a tight time frame regardless of the temporal context. This hypothesis was tested in Experiment 2.

## Experiment 2

This experiment sought to further assess the mechanisms of state transformation during memory maintenance. Building on the findings of previous experiment, we conjectured that, in the condition that the temporal context was beyond participants’ anticipation, the subsequent sensory stimuli were urgently processed during the state transformation process in case the sensory stimuli visually disappeared; this thus might result in an overlap and resources competition independently of the temporal context.

## Method

### Participants, stimuli and procedure

A sample of 30 participants (5 males) were recruited (mean age: 21.13 ± 2.39 years). Similar to the procedure and stimuli of preceding experiment (see Figure 1), this experiment nevertheless made a slight modification. The presentation time (0.2 s vs. 0.5 s) of memory array 2 were independent of the experimental block. The variable of presentation time and load was mixed within each block. The participants were instructed to perform six blocks, consisting of 192 trials in total.

## Result

Basing on the results of Experiment 1, we only analyzed the passive memory performance to test whether the active load had effect on the passive. An ANOVA with the variables of load (set size 2 vs. set size 4) and presentation time (200ms vs. 500 ms) was conducted for the passive memory accuracy (Figure 3). A significant main effect of load was detected (*F* (1, 29) = 16.335, *p* < 0.001, *η*^*2*^*p* = 0.360), but the interaction between them was not statistically significant (F < 1). The results suggested that passive representations were always affected by the active load variation when the temporal context was unpredictable.

**Figure 3:**
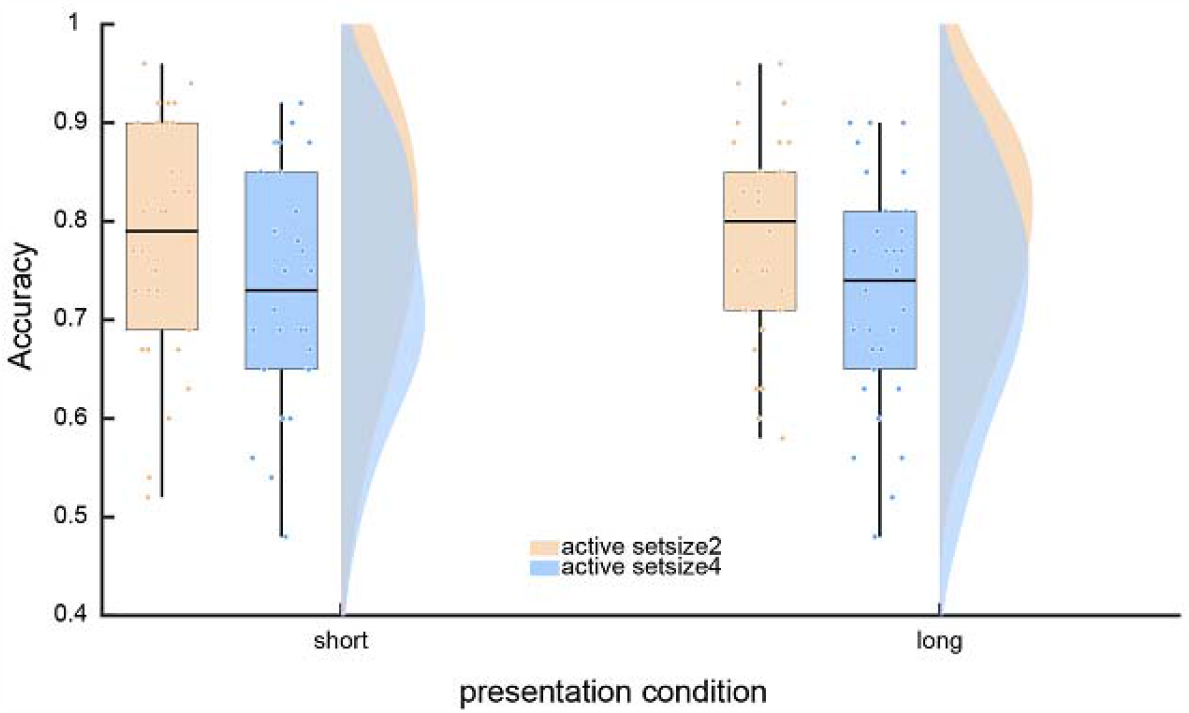
Mean accuracy of passive representations in Experiment 2. Boxplot, the middle lines indicate the median; the box outlines represent the 1^st^ and 3^rd^ quartiles. Scatters indicate data distribution for each condition.

## Discussion

As expected, participants initiated the subsequent sensory processing before the completion of state transformation though there was sufficient time for the two processes occurring sequentially, when the temporal context was unpredictable. Together with findings of Experiment 1, these findings consistently demonstrated that the state transformation mirrored a cognitive process that consolidated the memory representations into the passive state.

## Experiment 3

In previous two experiments, the to-be-transferred representations were not consolidated into the passive state during the delay retention. We then reckoned that the 800 ms delay retention was seemingly insufficient to allow for the completion of state transformation. Then we expected that the sate transformation could complete prior to the presentation of subsequent stimuli if extending the delay retention, thus never being interfered by new stimuli processing. Based on the experimental parameters of Experiment 1, we selected 800 ms vs. 1000 ms as the delay retention to test our hypothesis.

## Method

### Participants, stimuli and procedure

Another sample of 30 was recruited (5 males; mean age: 21.97 ± 2.70 years). This part repeated “stimuli and procedure” of Experiment 1 and made a slight modification (see Figure 1). Memory array 2 appeared for a fixed time of 0.2 s. While the delay interval between two memory arrays lasted 0.8 s or 1 s in a blocked design. The active set size 2 and 4 were randomly presented in each trial with equal probability. We designed 5 blocks of each delay condition, creating 320 trials totally for each participant. The order of short and long delay blocks was balanced between participants.

## Results

The collective data underwent the similar analysis plan to Experiment 1. A 2 (active set size 2 vs. active set size 4) × 2 (active state vs. passive state) × 2 (0.8 s vs. 1 s) ANOVA revealed a statistically significant interaction between representational state and active load, *F* (1, 29) = 45.259, *p* < 0.001, *η*^*2*^*p* = 0.609. Then we run a 2 × 2 ANOVA with factors active load and representational state for the short and long delay conditions separately. In the short delay condition (Figure 4A), the results showed a statistically significant interaction, *F* (1, 29) = 17.522, *p* < 0.001, *η*^*2*^*p* = 0.377. Subsequent test analysis suggested that the active memory accuracy was much higher in the set size 2 than the set size 4, *t*(29) = 6.749, *p* <0.001, Cohen’s *d* = 1.232, *BF10* = 75595.299. While the passive memory was also higher in the smaller active set size condition, *t*(29) = 3.044, *p* = 0.005, Cohen’s *d* = 0.556, *BF10* = 8.249. These results showed that the passive memory was affected by active load variation (i.e., an overlap of state transformation with subsequent sensory processing), which replicated the results of Experiment 1.

**Figure 4:**
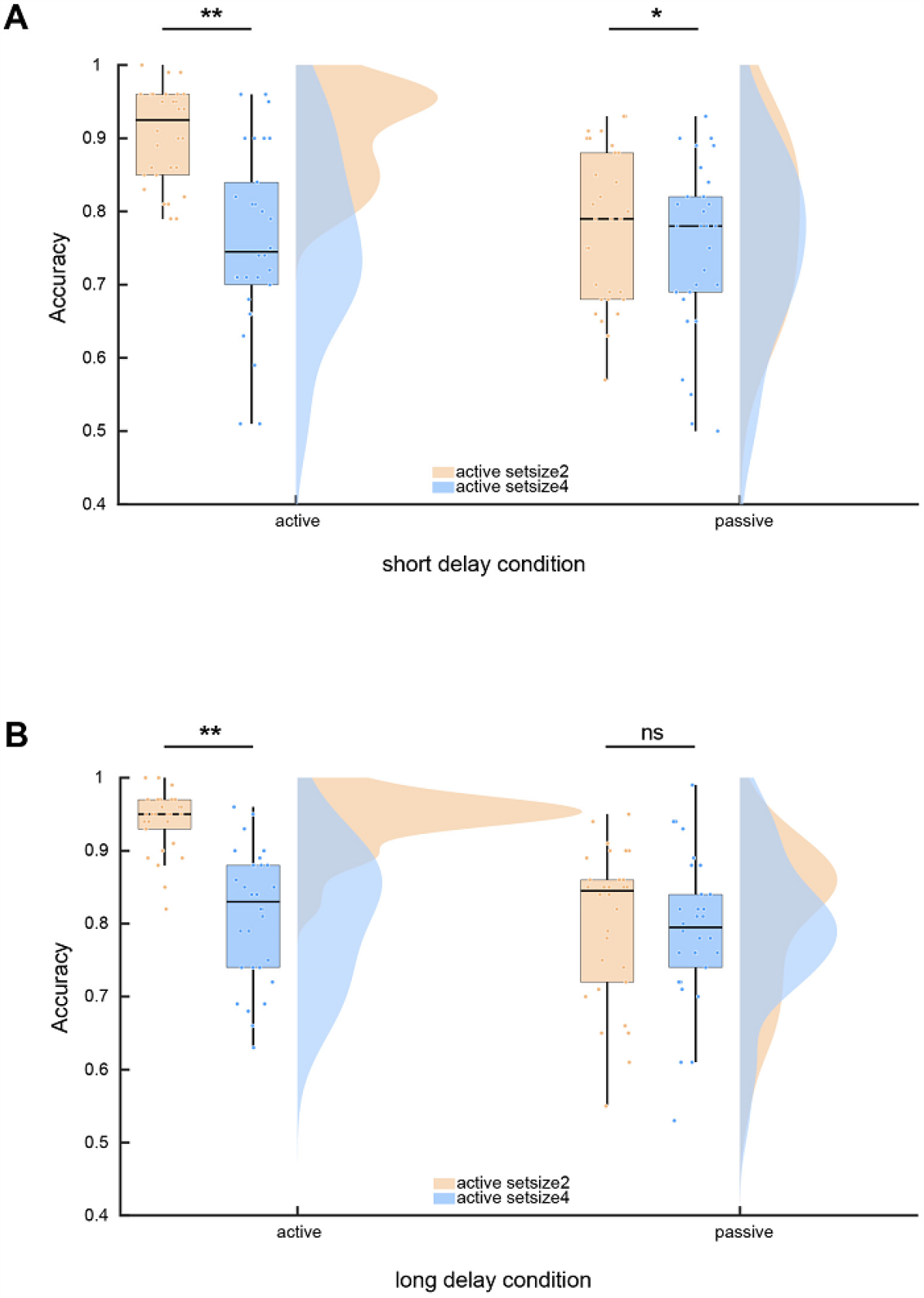
Mean accuracy of active representations and passive representations in short (A) and long (B) delay conditions in Experiment 3. Boxplot, the middle lines indicate the median; the box outlines represent the 1^st^ and 3^rd^ quartiles. Scatters indicate data distribution for each condition.

The analysis on the long delay condition was of us interest (Figure 4B). The 2 × 2 ANOVA results showed that the representational state significantly interacted with active load, *F* (1, 29) = 36.239, *p* < 0.001, *η*^*2*^*p* = 0.555, which was driven a null effect of active load on the passive memory. The subsequent test analysis showed that the active memory suffered a decrease when the active set size increased from two to four, *t*(29) = 8.682, *p* < 0.001, Cohen’s *d* = 1.585, *BF*10= 7.879e+6. Whereas the passive memory did not differ between the conditions of active set size 2 and 4, *t*(29) = 0.813, *p* = 0.423, Cohen’s *d* = 0.148, *BF*10 = 0.263. These results showed that extending the delay removed the active load effect on the passive representations.

## Discussion

In this part, these results indicated that extending the delay retention potentially conduced to a smooth state transformation. Such that the to-be-transferred items had been consolidated into the passive state before the presentation of subsequent stimuli, thereby never overlapping with the subsequent sensory processing. By contrast, due to the insufficient delay retention, the state transformation continued beyond the retention interval, and proceeded during the presentation of subsequent new input.

Overall, it could be known that the state transformation could accomplish the consolidation of passive representations during the delay retention as long as the delay retention was sufficient in time; this also indicated that transformation process was not triggered by the subsequent stimuli, but rather subject to the executive control.

## General discussion

The present study aimed to investigate how the state transformation of memory representations implemented during the multi-state maintenance. We adopted sequential presentation memory task, which effectively encouraged participants to offload the first array to the passive state when actively processing the subsequent stimuli. The passive representations should be independent of the active load variation according to the resource-dissociation account (Li et al., 2021), however, the active load effect on the passive representations was detected in some scenarios. Across three experiments, the behavioral results attributed the odd effect to an overlap between state transformation and subsequent sensory processing, due to the physically or psychologically insufficient temporal context. Accordingly, the current findings concluded that the state transformation reflected a consolidation process of memory representations into the passive state. A sufficient temporal context could lead to a smooth state transformation, thus avoiding the overlap and resource competition with subsequent processing. While the sufficient temporal lost benefit to the state transformation when the temporal context was beyond individuals’ anticipation.

Moreover, these results suggested that the state transformation could be completed either during the retention interval or proceeded to the presentation of subsequent stimuli; it follows that the state transformation process was cognitively controlled, not necessarily triggered by the appearance of subsequent new input.

The current results in fact fall in with dissociation account of VWM. Notably, the temporal context played a critical role in a smooth transformation of active representations into the passive. Previous studies have suggested that event processing that occurs in close proximity on a psychological timeline may lead to interference with each other (Shipstead & Engle, 2013; Souza & Oberauer, 2014; Brown et al., 2007). Thus, providing enough time for a smooth transformation prior to subsequent active processing contributed to a perfect dissociation between passive representations from the active processing. In terms of the resource competition in the relatively tight temporal context, we reasoned that the cognitive resources for the state transition was conceivably not domain-specific, but shared with sensory consolidation of new stimuli into the active state (LaRocque et al., 2013).

The current behavioral performance provided a new perspective to explicate the mechanisms of representational maintenance in the two distinct states. These findings revealed that, for a memory item encoded into VWM, the persistent neural activity underlying the memory maintenance in the active state did not accompany with the synaptic plasticity for transient maintenance in the passive state. In other words, the emergence of short-term synaptic plasticity for “activity-silent” pattern in the passive state came with the consolidation of memory representations into that state.

Considering that long-term memory was maintained as a pattern of the synaptic weights, transferring memory items into the passive state might render advantages to long-term memory performance for these items (McCabe, 2008; Rose et al., 2014), because state transformation promoted the formation of synaptic pattern in memory maintenance. The current findings about the nature of state transformation are indirectly obtained from behavioral performance; thus, the neural mechanism underlying the state transfer is necessarily explored with electrophysiological and neuroimaging techniques for obtaining more neurophysiological supports. Further research is needed to advance the understanding of state transformation in VWM.

In the current study, we proposed that the process of state transformation unfolded over time. This was consistent with the neural activity of EEG recordings which showed that the time required for memory representations to fall to the baseline level was roughly 1.25 s, and the time estimate for offloading the memory items into the passive state from behavioral studies (LaRocque et al., 2013; Oberauer, 2001, 2005). Though there was strong evidence for the proposal that sufficient time should be available for removing the currently task-irrelevant items into the passive state, that time span estimated blurred the time available for the state transformation itself. The current results indicated that a sufficient blank time between two arrays might afford the opportunity for state transformation, but may fail to ensure its completion. This motivated us to conclude that a perfect state transformation of memory items involved two kinds of time-related factors: one was sufficient blank retention that acted as the premise for state transformation (LaRocque et al., 2013; Li et al., 2021; Oberauer, 2001, 2005); the other was an appropriate time period reserved for consolidation of memory items into the passive state to avoid the resource competition with subsequent sensory processing.

The current research provided a further step toward to understanding the dynamic processing of VWM. Different from the recurrent neural activity, the synaptic changes in connectivity are far more energy-efficient since they do not require continuous sustained firing (Masse et al., 2020; Rose, 2020). Thus, the representational transition into the passive state might manifest great evolutionary advantage on some degree (Chota & van der Stigchel, 2021; Stokes, 2015). In the current study, the number of passive representations was fixed at two. That leaves up the question whether more time should be secured for a smooth transformation of more than two items. What’s more, these findings about how the state transformation works are obtained from behavioral results through three experiments; thus, the neural mechanisms underlying the state transformation is necessarily explored with electrophysiological and neuroimaging techniques. Further research needed to be conducted to advance the understanding of multistate maintenance of VWM.

## Conclusions

In summary, during the multi-state maintenance of memory representations, the state transformation acted as a process of consolidating memory representations into the passive state. The state transformation process could complete during the retention interval if it was sufficiently long; otherwise, the process continued during the presentation of new stimuli. We thus deduced that the state transformation was less related with the physical appearance of new stimuli, but driven by the need of subsequent cognitive processing. A smooth state transformation necessitated sufficient temporal context, thereby reducing memory cost and ensuring a behaviorally independence of working memory representations in different storage states.

## Acknowledgment

This research was supported by grants from the National Natural Science Foundation of China (NSFC31970989).

